# Per-ischemic changes in penumbral blood supply and its microscopic distribution

**DOI:** 10.1101/2023.06.07.544164

**Authors:** Nina K. Iversen, Eugenio Gutierréz Jimenéz, Peter Mondrup Rasmussen, Hugo Angelys, Irene Klærke Mikkelsen, Tristan R. Hollyer, Leif Østergaard

## Abstract

Acute ischemic stroke (AIS) is a frequent cause of death and adult disability. AIS patient management targets the *ischemic penumbra*: Hypoperfused, electrically silent brain tissue, which can be salvaged by restoring blood flow during the first, critical hours after symptom onset. Neuroimaging studies in AIS patients suggest that penumbral tissue is characterized not only by hypoperfusion, but also by microvascular flow disturbances that strongly affect tissue outcome. Here, we demonstrate that microvascular flows become increasingly chaotic in the ischemic penumbra in the hours after middle cerebral artery occlusion in a rat model of AIS. Biophysical models suggest that these disturbances are accompanied by increasing hypoxia in the absence of blood flow changes. Unlike findings in severe ischemia, pericyte constrictions do not appear to occlude penumbral capillaries. We propose that microvascular flow disturbances represent a critical feature of penumbral tissue, and a potential target for neuroprotective therapy after AIS.

## Introduction

Stroke is the second-most common cause of death, claiming 5.8 million lives worldwide in 2016^1^. Two out of three patients survive their stroke, but about 50% do so with permanent disabilities. As a result, stroke was also the second-most common cause of disability-adjusted life years lost to disease in 2016^2^. The majority of acute strokes are the result of focal brain ischemia caused by arterial occlusion, and the acute management of these patients therefore aims to limit brain infarction by restoring blood supply to the affected tissue^3^. The pathophysiological rationale for this therapeutic approach is the existence of an *ischemic penumbra*^4^: A volume of electrically silent, critically hypoperfused tissue, which can be salvaged by reperfusion within the first few, critical hours after symptom onset^5^. Penumbral tissue is further defined by its residual blood supply: while tissue injury ensues almost immediately at cerebral blood flow (CBF) below 8-12 mL/100mL/min^6^, brain tissue can survive for some time at CBF levels below the *ischemic threshold* of approximately 20mL/100mL/min^4,5^. Left untreated, the *ischemic core lesion* therefore grows as penumbral tissue dies, and penumbral contributions to the patient’s neurological symptoms become permanent. Over the past decade, intravenous administration of thrombolytic agents has been shown to improve functional outcome for patients when administered within 4.5 hours after symptom onset^2^ and in wake-up strokes^7^ in eligible patients, while endovascular clot-removal is proven safe and efficacious up to 24 hours after symptom onset^2^.

Several reports suggest that knowledge of CBF alone does not suffice to predict tissue viability after acute ischemic stroke. Perfusion magnetic resonance imaging (MRI) studies in acute stroke patients (< 12 hours after symptom onset) show that the survival of tissue with a certain level of hypoperfusion, as indexed by the prolongation of blood’s mean transit-time (MTT) through each image voxel’s vasculature, also depends on the within-voxel distribution of the blood flow^8-10^. Accordingly, tissue with an abnormally *narrow* within-voxel flow distribution on acute MRI shows extreme risk of subsequent infarction in the absence of reperfusion therapy – largely independent of its level of (hypo)perfusion^8-10^. Quantifying this phenomenon by the coefficient of variation (CoV) of within-voxel vascular transit times (their standard deviation divided by their mean, MTT)^11^, Engedal et al.^12^ found that low CoV in acute (< 6 hours after symptom onset) stroke patients is a common property of severely hypoperfused tissue (as measured by MTT), and associated with high risk of infarction if reperfusion cannot be achieved. Notably, in the absence of reperfusion, hypoperfused tissue with *normal* CoV shows much lower infarct risk than equally hypoperfused tissue with *low* CoV, confirming that not only the level of hypoperfusion, but also the extent of microvascular flow disturbances, impact the fate of hypoperfused tissue^12^. Reperfusion, meanwhile, reduces infarct risk of tissue with low initial CoV to the level of similarly hypoperfused tissue with normal CoV^12^.

The dependency of penumbral fate upon the microscopic distribution of residual blood flow raises an intriguing question: can hypoperfused tissue be protected by restoring or preserving bloods microvascular distribution – irrespective of whether subsequent reperfusion can be achieved? Penumbral microvascular flow disturbances might reflect reversible instances of phenomena observed during more severe ischemia, such as compression by swollen astrocytic end-feet, capillary obstructions by recruited immune cells, or constriction due to capillary pericyte rigor^13-18^. Indeed, if blood’s microvascular distribution could be preserved by therapies that are suitable for pre-hospital administration (See e.g., Gaudin et al.^19^), stroke patients might benefit irrespective of their access to or eligibility for subsequent reperfusion therapy.

MRI-based CoV-observations summarize the hemodynamics of thousands of interconnected microvascular segments within ~10 mm^3^ image voxels^11^. With respect to the nature of underlying microvascular flow disturbances, CoV estimates are therefore ambiguous, as they could reflect either compensatory homogenization of blood flows through *open* capillaries to optimize oxygenation as perfusion pressure drops^8,20^, gradual *cessation of flow* through microvascular pathways with long transit times^12^, or both. Although ischemia-related microvascular changes have been reported^18,21-24^, these studies have not, or to a limited degree, distinguished between core and penumbra tissue.

The aim of this study was therefore to characterize microvascular flow disturbances in penumbral tissue in a rat model of acute ischemic stroke and their evolution over the critical, first four hours after ischemia onset. First, we adapted the MRI CoV-method for dynamic two-photon microscopy (TPM) to examine the hemodynamics of the microvasculature that connects individual, diving arterioles and ascending veins in ischemic cortex. Next, we characterized the passage of erythrocytes through individual capillaries in the cortical microvasculature in order to characterize CoV changes in terms of the underlying hemodynamic and their severity. To this end, we applied biophysical models to estimate the impact of microcirculatory flow disturbances on penumbral oxygenation, and examined whether pericyte constrictions^18,25^ might affect capillary hemodynamics within penumbral tissue.

## Results

### Dissociated penumbral micro- and macrovascular response to middle cerebral artery occlusion

Following filament occlusion of the middle cerebral artery (MCAo) in Sprague-Dawley rats, we observed immediate reductions in the diameters of cortical pial arterioles (Fig. 1a,b) and in CBF as measured by the Laser Speckle Contrast Imaging (LSCI) signal over the affected cortical area (Fig.1c). We used the extent of LSCI signal reduction to subdivide hypoperfused cortex into penumbral, core, and control tissue, respectively^21-22,26-28^ (Fig. 1d), and verified the fate of penumbral tissue by TTC staining (Fig. 1e and Supplementary Fig. 1d).

**Figure 1.**
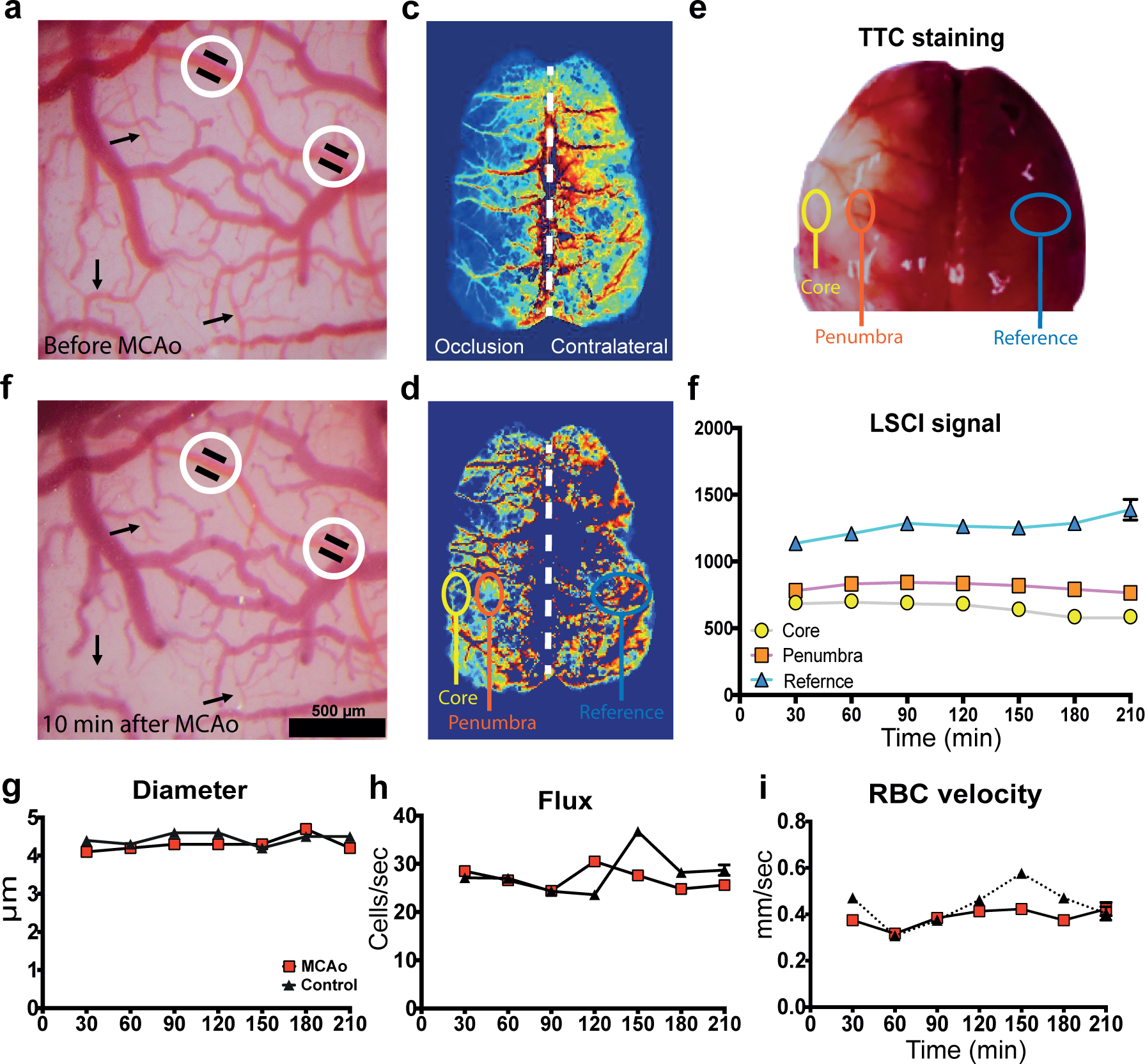
CBF and red blood cell kinetics in perfused capillaries after MCAO. Typical light microscope images of the rat cortical vasculature (a) before and (b) 10 minutes after middle cerebral occlusion (MCAO) in a rat. Black arrows indicate small arterial branches and cortical arterioles whose diameter decrease dramatically 10 minutes after occlusion. Within the white circles, the black lines indicate the outer arterial circumference before occlusion to visualize the diameter change. The scale bar indicates 200 μm. (c) Laser speckle contrast (LSC) image after 30 min of filament occlusion. Red colors correspond to high image intensity and high blood flow, whereas blue colors represent low LSC image intensity and low perfusion. (d) The same image thresholder so only tissue with image intensity between 25-50% of the contralateral hemisphere average is shown (the white dotted line indicates midline). Penumbral tissue was identified as tissue with LSC image intensity between 25 and 50% of contralateral tissue (pink) and infarct core (grey) as tissue with image intensity less than 25% LSC image intensity compared to a contralateral tissue. The core area was further verified by 2,3,5-Triphenyltetrazolium chloride (TTC) staining of the intact brain (e), where the white tissue shows the core 4h following MCA occlusion. The contralateral region of interest (turquoise) encircles the mirror-regions of the two, relative to the mid line (dotted while line). (f) LSC image intensity signal as a function of time for the core, penumbra, and contralateral mirror region. Asterisk indicates statistical significance between the three areas. (g) Comparing MACO and control animals, red blood cell (RBC) kinetics measured by two photon microscopy (TPM) line scanning of 530 individual penumbral capillaries (270 in control animals, 260 in MCAo, n= 8 for each group) every 30 min after filament occlusion provided estimates of capillary diameter, (h) RBC flux, and (i) RBC velocity. Asterisk indicates statistical significance (p < 0.0001) by two-way ANOVA test and Tukey multiple pairwise comparisons test.

Unlike the cortical LSCI signal (Fig. 1f), RBC velocities and flux (Fig. 1i,h), as measured by TPM line scans across 530 capillaries (270 in controls and 260 in stroke animals in the 100-250μm depth range), remained largely unaltered in penumbral micro vessels compared to those of control tissue. Also, capillary diameter did not change significantly (Fig. 1g), unlike pial artery diameter (Fig 1a,b). See Supplementary Material Fig. 2 for corresponding linear density and velocity variance measurements over time.

### Erythrocyte flows in penumbral capillaries deteriorate over time

Although average LSCI flow and capillary RBC fluxes remained largely constant in penumbral tissue for the duration of the experiment, RBC flow dynamics in individual capillaries grew increasingly chaotic over time. Observing RBCs within capillaries by two photon laser scanning microscopy (TPM) line-scanning (Fig. 2a), we observed instances of *flow reversal* in penumbral capillaries during 30 sec observation periods, as illustrated in figure 2b. Over the 4h observation period, the proportion of capillaries with flow reversal increased dramatically, ultimately involving as many as 25 % of the observed capillaries (Fig. 2d). This phenomenon was accompanied by a high incidence of *stalled flow*: capillaries with zero flux comprised 15-35% of the observed capillaries throughout the duration of the experiment (Fig. 2c). Capillary red blood cell (RBC) velocities were determined the angle resulting from their movements within FITC filled capillaries during TPM line scans (Fig. 2a). In addition to stalled flow and flow reversal, we observed increasing variability in penumbral RBC velocity when compared to control tissue. RBC variance and linear density are shown in Supplementary Fig. 2. We note that microvascular haematocrit was elevated in penumbral capillaries over the measurement period.

**Figure 2.**
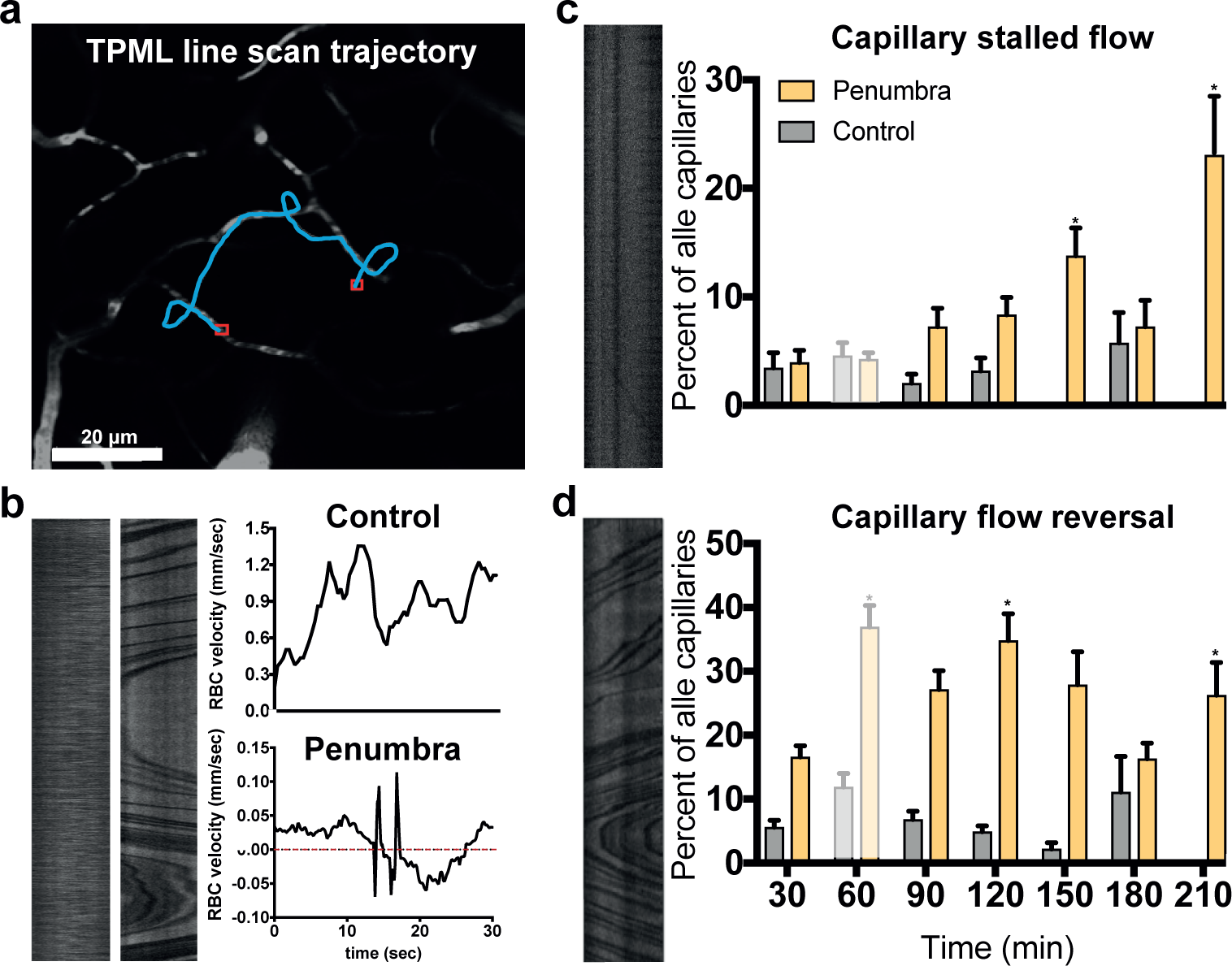
TPM line scans reveal increasingly chaotic RBC kinetics after MCAO. Using two-photon line scanning, 530 capillaries were scanned (270 in control animals, 260 in penumbral tissue of MCAo in the depth range between 100-250μm. Twenty-five capillaries were excluded from further analysis as the analysis yielded unrealistic cell densities (in excess of 300 cells/mm. (a) Typical line scanning trajectory. Injected FITC makes plasma appear bright while individual red blood cells (RBCs) can be observed as dark shadows within the vessel lumen. Two scan paths were prescribed for each capillary; one along the axis for RBC velocity estimation, and one transversal scan for RBC flux and capillary diameter assessment. (b) Typical line scan data for cortical capillaries in control animal and penumbral tissue of an MCAO animal, respectively. The corresponding, 30-second velocity profiles are shown for the control animal capillary (top) and a capillary within penumbral tissue (bottom). Note that RBC flow direction changes during the 30 second observation period (red dotted line indicates zero velocity). (c) shows the percentage of penumbral capillaries with such ‘chaotic’ behaviour as a function of the duration of ischemia. Note how the percentage of capillaries with reversing flows remains constant in normal tissue but increases dramatically in penumbral capillaries. (d) shows (left) the line scan of a capillary with stalled flow (flux=0) and (right) the percentage such capillaries represented of the total number of scanned capillaries for control and penumbral tissue over time. Asterisk denotes statistically significant difference (p<0.05) by two-way ANOVA test and Tukey multiple pairwise comparisons test. See Supplementary Material for further information about the 60 min data point.

### Progressive capillary transit time disturbances across penumbral tissue

To determine the course of hemodynamic disturbances in terms of the CoV of capillary transit times, we performed dynamic TPM of arteries and veins during bolus dye injection (Fig. 3) and conducted indicator dilution analysis equivalent to that of MRI studies to determine the mean and the standard deviation of plasma transit times as blood passes through the cortical microcirculation^30^ (Fig. 3d,e). Referring to the two as the mean transit time (MTT) and transit time heterogeneity (CTH), respectively, CTH is expected to change in proportion to MTT in normal, passive compliant microvascular networks^31^ such that their ratio, CoV, remains constant. Meanwhile, MTT is proportional to the dye’s vascular distribution volume (plasma) and inversely proportional to the plasma flow between arteriole and venule.

**Figure 3.**
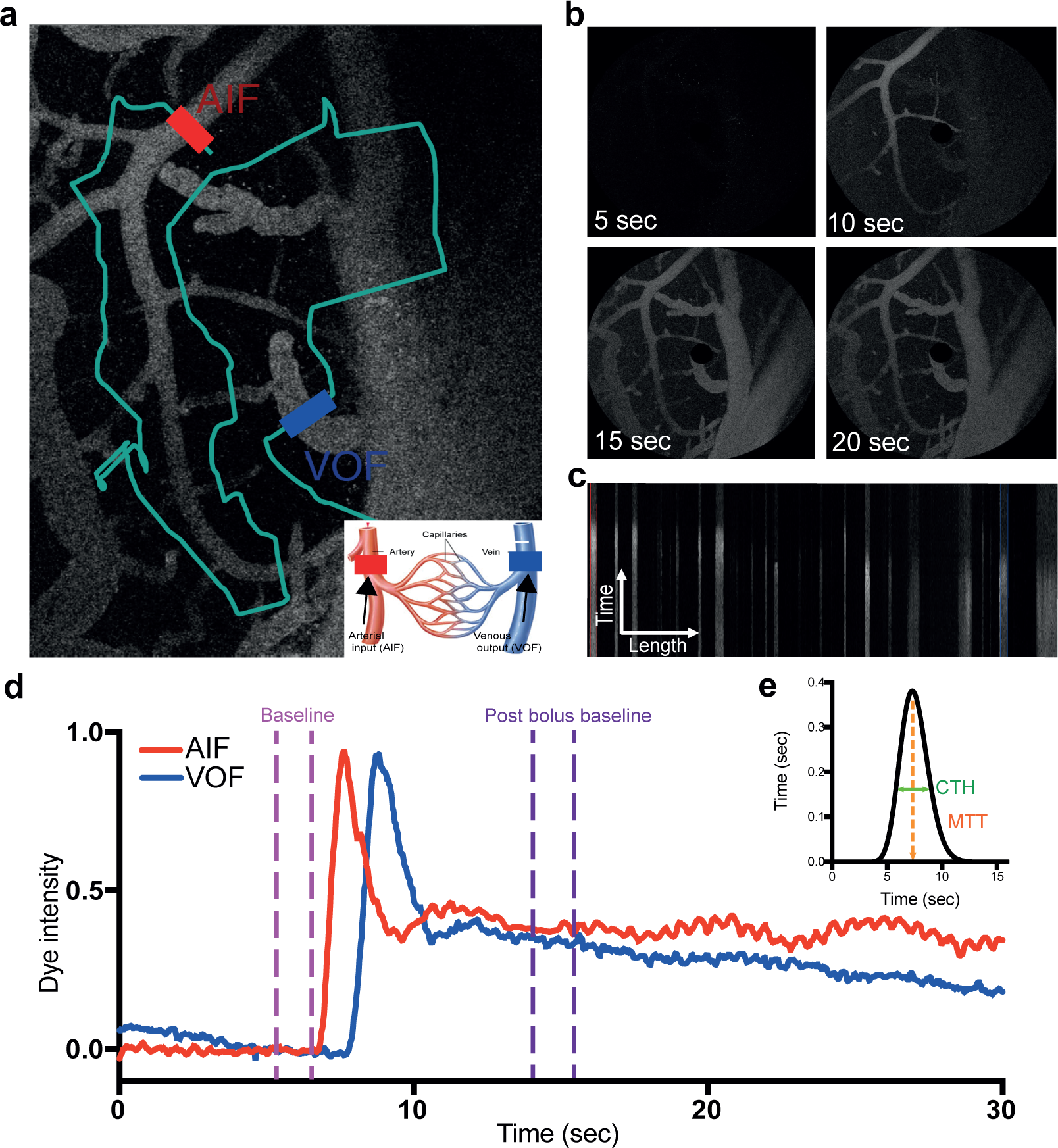
TPM bolus tracking technique in the penumbral area. A) Arterial input function (AIF) and venous output function (VOF) for quantifications of mean transit time (MTT) and capillary transit time heterogenity (CTH) in the capillary network. B). A total measurement of 303 bolus pairs were performed with 104 in controls and 199 in MCAo animals at a depth of 80-100 μm from the selected 0-position at the z-axe identified by a 20 sec spiral scan (512x512 pixels, time per pixel = 1.2 μsec.). The vessels were scanned as defined by the scanning route (grey line) crossing all AIF and VOF in the field of view (FOV). Red line marks the main AIF and blue the main VOF. The time series of the bolus injection is shown in C. Based on the scans, the 60 sec line scanned by time and length is shown in D), where the red and blue mark corresponds to the vessels marked in B. From the AIF and VOF intensity curves as a function on the 60 sec scan time, a 5 sec curve matching region was selected and curves were adjusted according to this including a baseline region and a post baseline. Scale bars = 200μm.

As expected, penumbral MTT values increased after vascular occlusion, reaching a peak at 120 min after which they decreased to values similar to those of control tissue after 210 minutes (Figure 4a). This biphasic behaviour was paralleled by penumbral CTH changes (figure 4b) such that penumbral CoV remained similar to control values until 180 minutes, after which penumbral CoV was significantly lower that of control tissue at 210 minutes (figure 4c).

**Figure 4.**
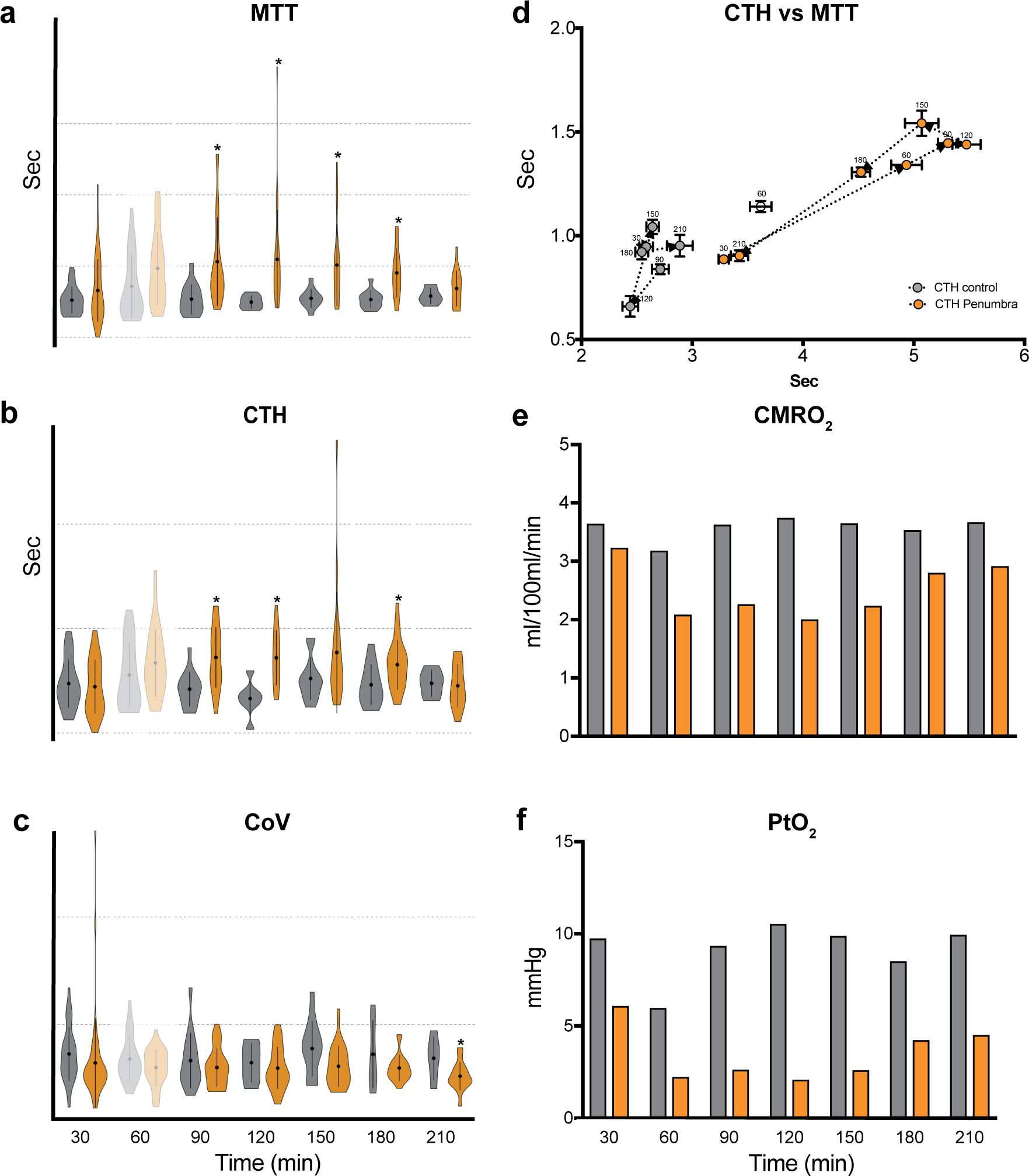
The transit time was disturbed across the penumbral capillary bed. Violin plots of mean transit time (MTT, a), capillary transit time heterogenity (CTH, b) and coefficient of variance (CoV, c) as a function of time every 30 min during the 4 h permanent occlusion period for both control (grey) and stroke (yellow) animals. d) CTH as a function of MTT for both control (grey circles) and stroke (yellow circles) visualized in an arrow diagram with the time of the bolus shown. Asterisk indicates statistical significance between groups (stroke, control) by mixed linear models. The data points from 60 min after occlusion (2^nd^ bolus) are faded out -see supplementary figure 1 for further information. The difference between groups at specific time points were tested by a t-test, p<0.01 Bar plots showing the estimated cerebral metabolic rate of oxygen (CMRO_2_, e) and tissue oxygen tension (P_t_O_2_, f) for a given MTT and CTH at each time point based on the models from {Angleys:2015iv}. Black bars indicate control animals, whereas yellow are MCAo.

We note that the apparent ‘normalization’ of MTT and CTH after 120 minutes happens as an increasing proportion of capillaries becomes affected by stalled flow and flow reversal. As penumbral blood flow, determined by the LSCI signal, changed little in this time period, we ascribe the decrease in MTT values to reduced vascular distribution volume for the injected dye. This volume is expected to decrease if the *diameter* or *total length* of passable microvascular paths from arteriole to venule decreases, or if microvascular haematocrit increases. Indeed, more capillary stalls further reduce the number of passable capillaries, just as increased haematocrit further reduces the dye distribution volume. Our measurements thus suggest that the decreases in MTT and CTH after 120 minutes represent a ‘pseudonormalization’. Simulations of microvascular flows suggest that gradual occlusion of capillaries tend to affect long pathways due to their higher resistance, giving rise to a reduction in CoV and a false impression of a ‘beneficial’ capillary flow homogenization^12^.

Thirty minutes after occlusion, an unexpected increase in blood flow in both ischemic and control animals was observed, resulting in elevated MTT and CTH values in both animal groups (Fig. 4a,b, Supplementary material Fig. 1b,c). These data are shown with transparent symbols in Figure 4, as it occurred in both MCAo animal and controls and seems to be a feature of our animal preparation, rather than of the stroke model.

### Metabolic significance of microvascular flow disturbances

To estimate the severity of the microvascular blood flow disturbances, the impact of the measured MTT and CTH changes on tissue oxygenation were calculated *post hoc*, using biophysical models^32,33^ to calculate corresponding, steady state cerebral metabolic rate of oxygen (CMRO_2_), and tissue oxygen tension (P_t_O_2_, Fig. 6). Note how CMRO_2_ and P_t_O_2_ in control animals remained constant throughout the experiment and within the normal range (CMRO_2_ =3.6-3.8 mL/100mL/min; P_t_O_2_ = 10-15 mmHg). In penumbral tissue, however, CMRO_2_ and P_t_O_2_, decreases to reach 1.98 mL/100mL/min and 2 mmHg, respectively.

**Figure 5.**
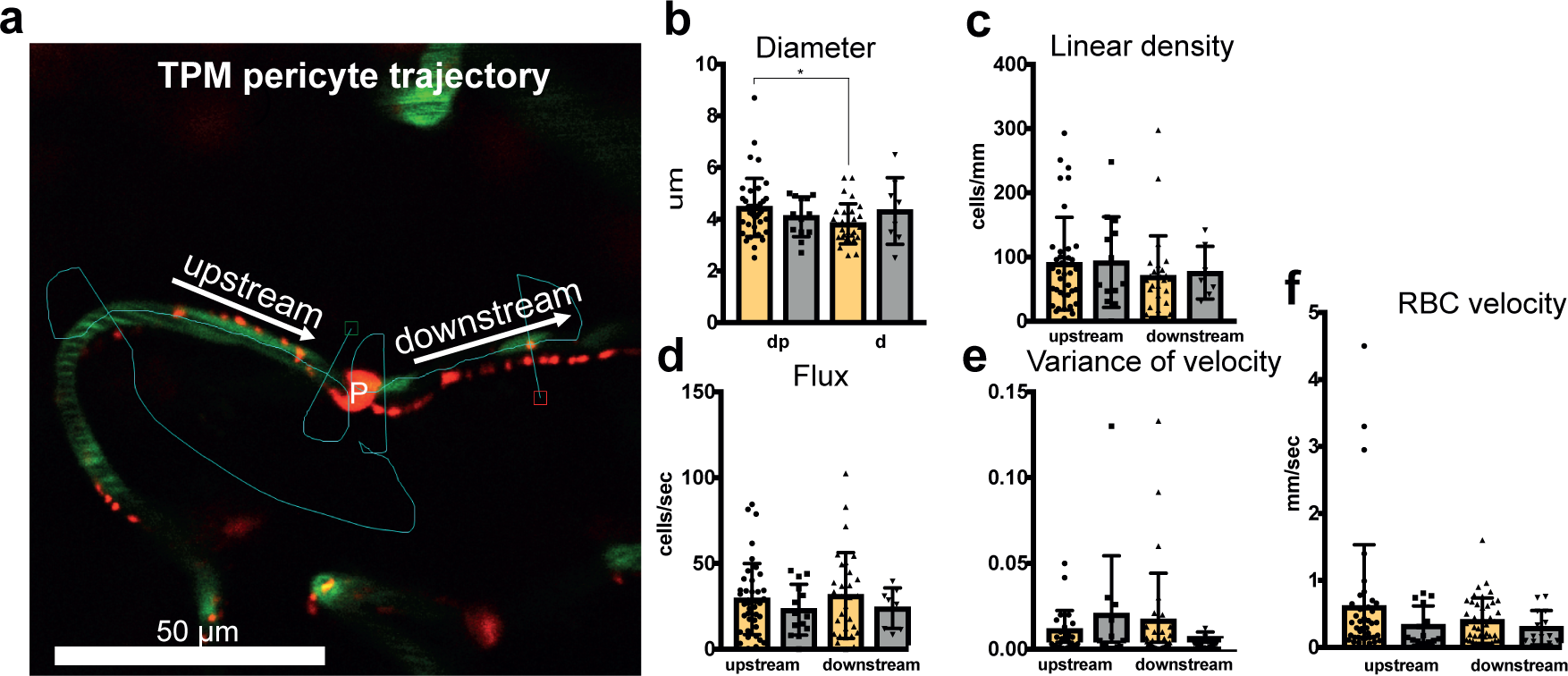
Capillary pericytes do not affect penumbral RBC hemodynamics *in vivo*. Two-photon image of a line scan of an individual capillary defined as a vessel showing single cell passages (cell’ shadows) within the vessel lumen. Pericytes in the penumbral region were stained by topical injections of Nissl bodies and were visible down to a depth of 350μm in the in both control and MCAo animals. They lined capillaries with multiple processes extending along vessel branches. The labelling was bright and concentrated at both cell soma and throughout the processes, where it displayed a punctate pattern. Two scan paths were performed on each capillary, along the axis for RBCv estimation, and a transversal scan for RBCflux and diameter assessment. Green colours denote plasma, whereas red colours represent a capillary pericyte cell body and arms. Line scans were made on both side of the pericyte and diameter scans were made next to the pericyte body and at a far distance, where 1 denotes upstream pericytes and 2 downstream (a). Data is shown in bar charts, where diameter at the pericyte cell body (dp) and at a far distance (p, b), linear density (c), RBC flux (d), variance of the line scan velocity profile (e) and RBC velocity (f). Black bars are control and yellow stroke. Asterisk indicates significance difference between upstream and downstream the pericyte in the specific type, i.e. control or stroke, p<0.05.

**Figure 6.**
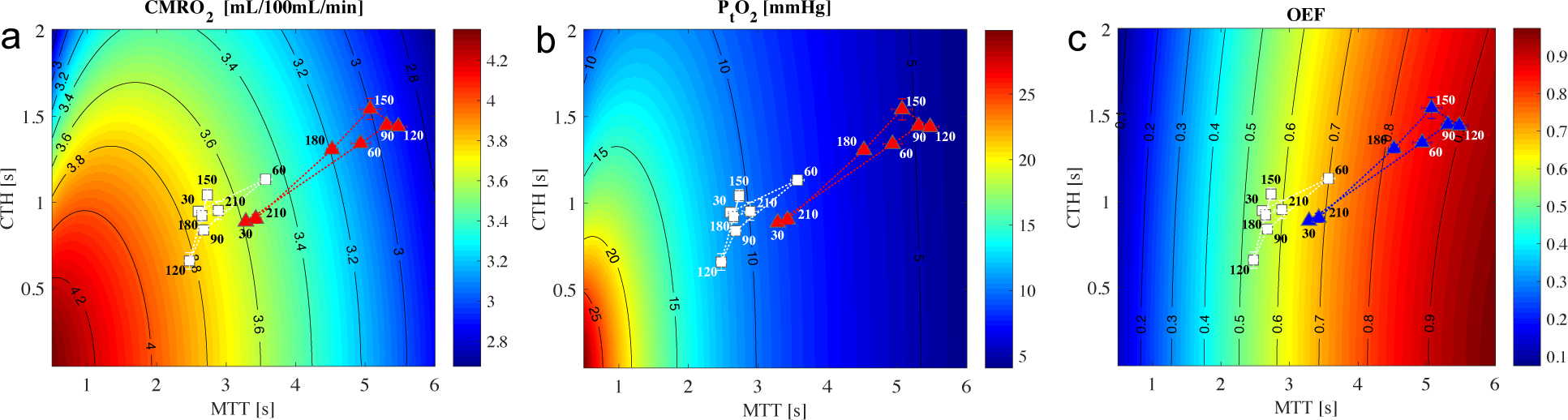
Contour plots reveals low penumbral tissue oxygen tension. To access the effects of capillary flow pattern changes on oxygen availability during stroke, cerebral metabolic rate of oxygen (CMRO_2_), tissue oxygen tension (P_t_O_2_) and oxygen extraction fraction (OEF) were predicted based on the MTT and CTH values in a biophysical model for oxygen extraction^32^ assuming all vessels stay open. The effect of increased CTH on tissue oxygen uptake i based on the theory by Jespersen and Østergaard^31^. According to the model, residual metabolism results in a low tissue oxygen tension (P_t_O_2_), and thus an elevated oxygen extraction fraction (OEF) until the network collapses, keeping in mind that the model assumes that vascular volume is preserved (all vessels are open). CTH as a function of MTT for both control (white boxes) and stroke (red/blue triangles) visualized in an arrow diagram with the time of the bolus shown.

### Capillary pericytes do not affect penumbral RBC hemodynamics

To examine whether hemodynamic changes were affected by changes in pericyte morphology, we performed TPM line scanning of individual capillaries, up- and downstream of pericyte soma, respectively, and measured capillary diameter at the location of the pericyte soma (fig. 5). Pericyte staining revealed a uniform distribution of these cells throughout the tissue volume, localized at capillary branch points as well as along individual capillaries. We found the inner diameter of the capillary to be significantly larger at the position of the pericyte soma than further away (Fig. 5b). RBC linear density was modulated in the same way by the soma passage in MCAo and control animals, while RBC flux was unaffected, confirming that no branching occurred at the location of the pericytes. In MCAo animals, the variance of RBCv was elevated upstream of the pericyte, which we ascribe to the increased, general level of flow disturbances across the capillary network, including capillaries with reversed flow (Fig. 5e). Accordingly, penumbral pericytes do not seem to explain trapped erythrocytes or stalled blood flow in penumbral capillaries (Fig. 2).

## Discussion

Our first main finding is that, during the first few hours following filament-induced MCAo in rats, microvascular flow disturbances evolve in penumbral tissue despite constant, residual blood flow. Thus, a growing fraction of capillaries showed reversal or complete cessation of RBC flows, particularly beyond 90 minutes of ischemia. Similar observations of the cortical microvasculature in murine experimental models during stroke-like conditions have revealed reduced capillary perfusion levels and pronounced microvascular flow disturbances^24^ in concert with irreversible pericyte constrictions^15,18,22,34,34^. Our study extends these by using established LSCI thresholds to distinguish between ischemic core and penumbral tissue, and by following the progression of hemodynamic changes over time^14^. With this distinction, our findings extend previous studies by ascribing microvascular flow disturbances in penumbral tissue to mechanisms other than pericyte constrictions.

Our second main finding is that, according to our biophysical models, oxygen availability dwindles over time in the ischemic penumbra, despite constant residual blood flow. Tissue oxygenation is traditionally gleaned from residual CBF, factoring in the higher oxygen extraction fraction that is characteristic of ischemic tissue. Therefore, current stroke management focuses on restoring blood flow as soon as possible. In parallel, neuroprotective strategies have sought to reduce the vulnerability of neurons to the corresponding, low oxygen levels, although translation of successful drug candidates to humans have failed. Due to the poorer oxygen extraction that results as this residual blood flow has to pass through fewer capillaries, however, oxygen availability deteriorates as a result of these microvascular flow disturbances. Indeed, our calculations suggest that normal oxygen metabolism cannot be sustained for more than 60 minutes of ischemia with deteriorating microvascular flows, according to our biophysical models. Together, our findings thus support the notion that penumbral oxygenation availability depends on both CBF and bloods microvascular, and that failing capillary flows contribute to penumbral infarction. Future studies should therefore examine the neuroprotective effects of therapies that support sustained capillary perfusion during ischemia.

The role of pericytes in capillary flow disturbances during ischemia remains unclear. Studies by Hall et al.^25^, Yemisci et al^18^. and Zhang et al^34^. show that pericytes die and constrict during ischemia, closing RBC flow through individual capillaries. It hould be kept in mind, however, that pericyte tone is affected by multiple factors^35^ that may change during whole animal- and subsequent tissue preparation. We searched for signs of altered pericyte morphology and function by examining whether pericytes located along capillaries in penumbral tissue might affect capillary diameter (Fig. 5b) or RBC passages, as inferred from the relation between upstream and downstream RBC velocities compared to control animals (Fig. 5f) *in vivo*, but found no clear indications thereof.

Potential causes of capillary flow disturbances include capillary occlusions or more proximal constrictions. Recent studies by Erdener^14,36^ found that dynamic stalling erythrocytes in capillaries occurred persistently in salvageable tissue (penumbra) after reperfusion and further that leukocytes were traveling slowly through capillary lumen or were stuck. Further, a decreased number of stalls were associated with improvement in penumbral blood flow within 2-24h after reperfusion along with increased capillary oxygenation and hereby improved functional outcome^36^. In addition, neutrophils adhering to distal capillary segments has shown to obstruct of 20-30% of capillaries in the infarct core and penumbra^37^. Grubb et al.^38^ recently showed the presence of a precapillary sphincter at the transition between the penetrating arteriole and the first capillary that links blood flow in capillaries to the arteriolar inflow and further that global ischemia and cortical spreading depolarization constrict sphincters and cause vascular trapping of blood cells. These results reveal precapillary sphincters as bottlenecks for brain capillary blood flow. We further speculate that capillary flow disturbances relate to swelling of astrocytic endfeet^39^, as we recently found in in a murine model of subarachnoid hemorrhage^40^.

### Limitations to the study

While we used LSCI to assure that our hemodynamic observations were performed in tissue that was initially penumbral tissue, our experimental design did not allow us to establish when in the course of the four-hour observation period tissue succumbed to irreversible tissue injury.

We see an increased degree of disturbed microvascular flow with gradually more irreversible damage already within the first 90 min of occlusion, which in concert with stalling and reversing RBC flows in individual capillaries led to highly elevated CTH. When the ischemic episode is initiated, the oxygen demands from the compromised penumbra tissue is met by increasing flow at 30 min (1^st^ bolus, Fig. 4d) illustrated by the increased MTT. We speculate that beyond 30 min of ischemic stroke, the further increased flow cannot continue to match the oxygen demand from the tissue as the flow pattern becomes more heterogeneous (increased CTH), and further that at 120 min after the stroke onset, the blood flow cannot sustain the oxygen delivery and the capillary network collapses. Future studies should reveal the relation between deteriorating capillary flows and these penumbral no-flow capillaries.

Our indicator dilution technique relies on repeated dye injections with parallel exposure of the microcirculation to laser light (Fig. 4), which might damage individual capillaries. Our repeated measurements in control animals show that these injections, and the vascular exposures to laser light, do not affect the hemodynamic responses and the estimated parameters. Accordingly, the measured parameters (MTT, CTH, Fig. 4a,b) remain stable during the whole experimental period with 7 injected boluses.

Our experimental approach did not allow us to study individual capillaries immediately after ischemia onset, and we would therefore overlook vessels that were closed to plasma flow as we commenced two-photon imaging. Thus, from our light microscope images (Fig. 1a), we found contracting capillaries 10 min after MCA occlusion, and not all of these were filled with the FITC dextran dye. Such capillaries would not appear in the TPM scanning.

## Conclusion

Our data provide direct *in vivo* evidence that the ischemic penumbra is characterized by deteriorating microvascular hemodynamics, as much as a low but constant, residual CBF. According to our biophysical models, these progressive, microvascular flow disturbances may be major contributors to the demise of penumbral tissue (infarction), consistent with earlier MR data. Our data do not support capillary pericytes as culprits in the disturbances. Further studies should directly quantify the effects of altered capillary blood flow patterns on penumbral oxygen availability and cell survival, and demonstrate that pharmacological improvements of capillary blood flow save penumbral tissue.

[Methods 3000 ord - lige nu: 2286]

## Methods

### Animals

Thirty-eight male Sprague Dawley rats (Taconic Biosciences, Denmark, mean w=380±1.7g) were studied as part of the experiments. All rats were $housed in standard cages in groups of two and kept on a 12-hour light:dark cycle with free access to food and water. All experiments were carried out in accordance with the regulations of the Danish Ministry of Justice and Animal Protection Committees. The study was approved by the Danish Animal Inspectorate with licenses no 2013-15-2934-00788 and 2018-15-0201-01443, and complies with the ARRIVE guidelines 2009 (Animal Research: Reporting In Vivo Experiments).

### Experimental protocol

The animals were studied according to one out of three protocols to address the following: Group I. Identification of the penumbra (w=334±3.3g, control n=2, stroke n=6); Group II. Capillary blood flow patterns (w=348±3.3g, control n=7, stroke n=8, Fig 1); and Group III Capillary RBC velocity (RBCv) and effects of pericytes (w=434±3.5g, control n=7, stoke n=8). In all groups, aninmals were randomly assigned to either the stroke or the control group by coin flip. None of the animals were used in more than one experimental protocol. The protocol lasted 4h and measurements were performed either continuously or every 30 min throughout this period.

### Surgical preparation and monitoring

The rats were intubated and artificially ventilated with 30% O_2_ using a small animal ventilator (Harvard Apparatus 683, MA, USA) under Hypnorm-Dormicum anaesthesia (1.8 ml/kg for induction and 1/3 dose every 30 min for maintaining). End-tidal CO_2_ was continuously monitored (MicroCapStar End-tidal CO_2_ analyzer, Cwe Inc., Ardmore, USA) and ventilation rate was adjusted to ensure end-tidal CO_2_ levels between 35-40 mmHg. Body temperature was maintained at 37°C by a heating pad controlled by feedback from a rectal thermometer (Homeothermic monitor, Harvard Apparatus, MA, USA). The right femoral artery was cannulated for continuous recording of mean arterial blood pressure (MAP) and heart rate (*f*_H_) using a BP-1 system (WPI Inc., Sarasota, FL, USA) and for monitoring of arterial blood pH, P_a_CO_2_, and S_a_O_2_ (ABL90 Flex, Radiometer Medical ApS, Brønshøj, Denmark). Three blood samples (60μL) was taking from each animal; before the MCAo, just after the occlusion and at the end of the experiment (Fig. 1). The femoral vein was cannulated in animals in experimental groups II and III for intravenous administration of fluorescence dyes. All physiological data were sampled at 50Hz with a Powerlab 35 series 16 bit data acquisition system (ADInstuments Ltd, Oxford, United Kingdom).

### Middle cerebral artery occlusion (MCAo)

Permanent focal cerebral ischemia was achieved by occluding the distal portion of the middle cerebral artery (MCA) as previously described by Belayev^41^ and Koizumi^42^, using an intraluminal thread and a 4-0 reusable suture (4041PK5Re, Doccol Corporation, Sharion, USA.). In brief, the carotid bifurcation was exposed and the common carotid artery occluded by a suture. The branches of the external carotid artery were dissected and divided, and the filament were then inserted into the common carotid artery and advanced 20 mm. The filament remained in place during the 4h experimental protocol to achieve a permanent MCA occlusion.

### Laser Speckle Contrast imaging

For penumbra identification in experimental group I, rats were placed in a stereotactic frame (Small animal stereotaxic, Kopf Instruments, CA, USA) with ear bars. The skull was thinned by a dental drill (Foredom Electrics CO, CT, USA) under continuous cooling with room temperature saline until the pial vessels became visible. The thinning procedure was done immediately prior to the MCA occlusion (MCAo). To increase the signal from the Laser Speckle Contrast Imaging (Moore Instruments, Devon, UK) and to keep the bone moisturised, almond oil was continuously added. Perfusion was measured continuously during the 4h experimental period after MCAo using a full-field laser perfusion imager (MoorFLPI, Moor Instruments Ltd., Devon, UK). A 785 nm class 1 laser diode was employed for illumination of the tissue down to a depth of approximately 1 mm. Laser speckle images was acquired using a 576 x 768 pixel grayscale CCD camera operating at a frame rate of 25 Hz. After acquisition, images were converted to matlab files and analysed using custom written matlab software.

### Cranial window preparation and pericyte labelling

For Two-Photon microscopy (TPM) studies, a chronic cranial window was established immediately prior to the MCA occlusion at the calculated expected position of the penumbra as determined from the laser speckle data above. A metal holding bar was glued to the right frontal bone of the rat to immobilize its head during imaging and a cranial window with 5 mm diameter was drilled through the parietal bone. Drilling was performed during continuous cooling by room temperature saline. After the skull was removed, the dura was peeled off carefully to avoid any bleedings. In order to visualise cortical pericytes, a topical application of Nissl bodies (NeuroTrace 500/525) was performed (Damisah et al., Nature Neuroscience, 2017) repeatedly during 10 min. Afterwards the window was cleaned by saline before being filled with 1.5 % agarose in saline (Sigma-Aldrich, Søborg, Denmark) and covered with a glass coverslip secured with dental acrylic (GC Fuji PLUS dental cement, GC Corporation, Tokyo, Japan). The window was allowed to dry for 45 min before subjecting animals to MCAo prior to TPM.

### Two-photon microscopy (TPM)

Imaging was performed using a Praire Ultima-IV In Vivo Laser Scanning Microscope (Bruker Corporation, Billerica, MA, USA). The field of view (FOV) was centred in the cranial window over the penumbra. To characterise the microvascular blood flow changes during the infarction of the penumbra, an indicator dilution technique was applied to determine the distribution of transit times through the cortical vasculature in experimental group II. This technique comprises dynamic imaging of the passage of a bolus-injected dye as previously described by Gutierrez et al^29^. Briefly, a 10X water immersion objective (Olympus 0.30 numerical aperture-NA, 3.3 mm working distance-WD) was used with a pixel resolution of 1.16-2.23μm per pixel (1x-2x) depending on the optical zoom. To adjust the FOV and set the coordinates, shadows originating from the pial vessels vessel were visualized by NADH auto-fluorescence using second harmonics (laser λ=810nm, optical parametric oscillator (OPO, λ=1100). A 50μl bolus of 0.5% Texas-red dextran solution (70,000 MW, 5 mg/mL in 0.9% NaCl, t_1/2_ ∼ 25min, Termo Fisher Scientific) was injected at a constant rate of 30μl/sec using a syringe infusing pump (GenieTouch, Kent-Scientific, Torrington, CT, USA) while performing a 20 sec spiral-scan at 6.25fps within a single plane (512x512 pixels, dwell time per pixel=1.2μsec). Fluorescent emission was detected by a GaAsP-PMT (Hamamatsu, H7422-40) using a 660/40 nm-emission filter to optimize signal to noise ratio. Laser λ=810nm, OPO λ=1100nm.

Arteries and veins were identified based on the timing of dye arrival (Fig. 4a). After identification of vessels within the FOV, a scan path was defined though the pial vessels of the upper cortically layer by applying the TPMs free-hand drawing tool (PraireView, Bruker Corporation, Billerica, MA, USA). Scanning lines were drawn to cross the largest artery and vein within the FOV as well as arterioles and venules. Subsequently, repeated line-scans were performed (6.7 msec per line-scan path dwell-time per pixel 1.2μsec) for a total scan time of 60 sec, during which the dye was injected after 10 sec. Bolus passage measurements were repeated every 30min for the entire 4h experimental period. To obtain an angiogram that displays the anatomical relation of the vessels, a z-stack was acquired on a FOV of 1.18μm^2^ (10x objective) from the pial surface down to a depth of 300-400 μm.

In order to characterise capillary RBC velocity (RBCv) and effects of pericytes in experimental group III, single capillaries were scanned to access RBCv and RBC flux (RBCflux) with a 20X water immersion objective (Olympus, 1.0 NA 2.0 mm) with a pixel resolution between 0.19-0.23μm per pixel depending on the optical zoom used (5x-6x). A single bolus of 50μL 0.5% Texas-red dextran solution (70,000 MW, 5 mg/mL in 0.9% NaCl, t_1/2_ ∼ 25min, Termo Fisher Scientific) was injected, and capillaries were defined as vessels showing single cell passages within the vessel lumen. Two scan paths were performed on each capillary, along the axis for RBCv estimation, and a transversal scan for RBCflux and diameter assessment. This was done on both side of the position of pericyte, and diameter scans were taken just next to the pericyte body and at the far end of the capillary. Laser λ=810nm, OPO=1100nm, Laser for pericyte λ=1000nm. Four sub-FOV at depths of 80-150μm was chosen within the main FOV and all capillaries in each sub-FOV (6-10) were scanned every 30 min during the 4h experimental period. Individual capillaries were scanned for 30 sec in the depth of 80-150μm.

### Laser Speckle image analysis and penumbra identification

The spatial extend of the ischemic penumbra was identified from the laser speckle images. First brain pixels were isolated by manually outlining brain pixels and a centre line was defined to separate the control and stroke hemispheres (Fig. 1d, Supplementary materials Fig. 1). Larger vessels had the highest speckle contrast and the corresponding pixels were masked out by applying intensity thresholds. Laser speckle has previously been used to visualise CBF changes throughout the ischemic territory in stroke models in rodents and cats^22,26,28,43^. Based on this approach, penumbral regions were defined by the group of pixels with intensities within the interval 25%-50% of the maximum intensity of the control hemisphere^33,44^ (fig. 1d,e). Two ROIs were drawn to represent the penumbra and the core within the stroke hemisphere, and these regions were mirrored along the center line to define corresponding ROIs in the control hemisphere. Finally, pixel intensities were averaged within the ROIs (control ROIs pooled, Fig. 1d).

### Bolus tracking image analysis

To identify the primary inputs and outputs for the bolus tracking analysis within each FOV, the vessels were separated according to their diameter and the time-to-peak (TTP) of their concentration curve (CTC). The arterial input function (AIF) was the CTC with a large diameter, which was first to enhance (i.e. had the shortest TTP, Fig. 3b), whereas the venous output function (VOF) was identified as the CTC of a large vein, which was the last to enhance (i.e. longest TTP). Then, diving arterioles and ascending venules were selected (Fig. 3a). From the z-stack scan, vessels were then paired based on their apparent, anatomical connectedness throughout the capillary bed. The dye transport was analysed based on these same pairs every 30 min. The pre-bolus intensity signal was identified for each vessel and the baseline signal was subtracted to create curves proportional to the dye concentration (CTCs). Afterwards, all curves were scaled to the post-bolus level of the AIF to correct for differences in signal intensity due to light traveling from vessels of varying depth (Fig. 3d). Analogous to the approach used when fitting the residue function-based tracer retention observed by DSC-MRI, we determined the transport function by deconvolution of the VOF with the AIF^11^. The resulting transport function comprises of a gamma cumulative distribution with two parameters α and β, from which the mean transit time (MTT) and capillary transit-time heterogeneity (CTH) were estimated as the mean (αβ) and the standard deviation 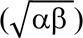 of the distribution, respectively. The coefficient of variation (CoV) of transit times was determined as the ratio between CTH and MTT 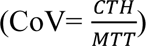. The deconvolution results were discarded if α<1 and β>0.5

### Capillary line scan and pericyte imaging analysis

RBCv values were calculated based on the Radon transformation^46^ as previously described in Gutierrez et al^29^. Flux was estimated by analysing the intensity variations occurring within the cross-sectional scan of each capillary. As for the velocity estimation, average intensity profiles were derived from 150 msec time intervals within the transversal line scan. Low and high contrast were taken to indicate presence and absence of RBC, respectively. The number of capillaries showing either stalled blood flow (flux=0) and/or reversed flow, where RBCv alternates between positive and negative flow values, was calculated. Linear density (LD) was calculated as flux/velocity and capillaries with a LD higher than 300cells/mm were discarded from further analysis since this represents an unrealistic high cell/plasma ration for an erythrocyte of approximately 2μm. Estimates of capillary RBCv and RBCflux were averaged for the 30 sec line scan time, and their standard deviation during the 30 sec period was calculated as a measure of the variance of flow.

Capillary diameter was estimated based on dye concentration line profiles from the line scan data. The vessel diameters were estimated as full-width at half maximum (FWHM) values of the line profiles as described earlier^29,46^, assuming the vessel intersects define their full diameters. Vessel cross-sectional areas were estimated from these diameter calculations. Vessels showing inner diameter ≥ 10μm were considered to be arterioles or venules and therefore disregarded in further analysis.

### Post hoc analysis

To access the effects of capillary flow pattern changes on oxygen availability during stroke, cerebral metabolic rate of oxygen (CMRO_2_), tissue oxygen tension (P_t_O_2_) and oxygen extraction fraction (OEF) were predicted based on the MTT and CTH values in a biophysical model for oxygen extraction^33^ assuming all vessels stay open (Fig. 6).

### Statistical analysis

Statistical analysis was performed in RStudio (Verision 1.1.456 © 2009-2018, RStudio Inc.) using a statistical significance level of 0.05. Physiological data were analysed by One-way and Two-way ANOVAs and Tukey multiple comparisons test. Differences in the hemodynamic values were tested by linear mixed model (LMM), where the hemodynamic values were treated as fixed effects whereas time and type (i.e. control vs stroke) were random effects. P-values were obtained by likelihood ratio test of the effect in questions. Effects of types at the specific time points was evaluated by t-test. The difference between the hemodynamic upstream and downstream of the pericytes, as well as the fraction of capillaries with stalled or reversed flows, were addressed by a two-sided t-test. All results are presented as their mean value ± SE, if not otherwise indicated.

## Acknowledgements

This study was funded by The Danish Research Council individual postdoctoral grant and Sapere Aude Young Research Talent (NKI).

## Author contributions

NKI designed the study, performed the experiment, analyzed the data and wrote the manuscript. EGJ contributed to the TPM scannings and data analysis. PMR and IKM wrote the matlab codes and contributed to data analysis. HA conducted the oxygen calculations, TRH contributed to the experiment and animal models. LØ contributed to the experimental design, data analysis and helped write the manuscript.

---

**Supplementary figure 1.**
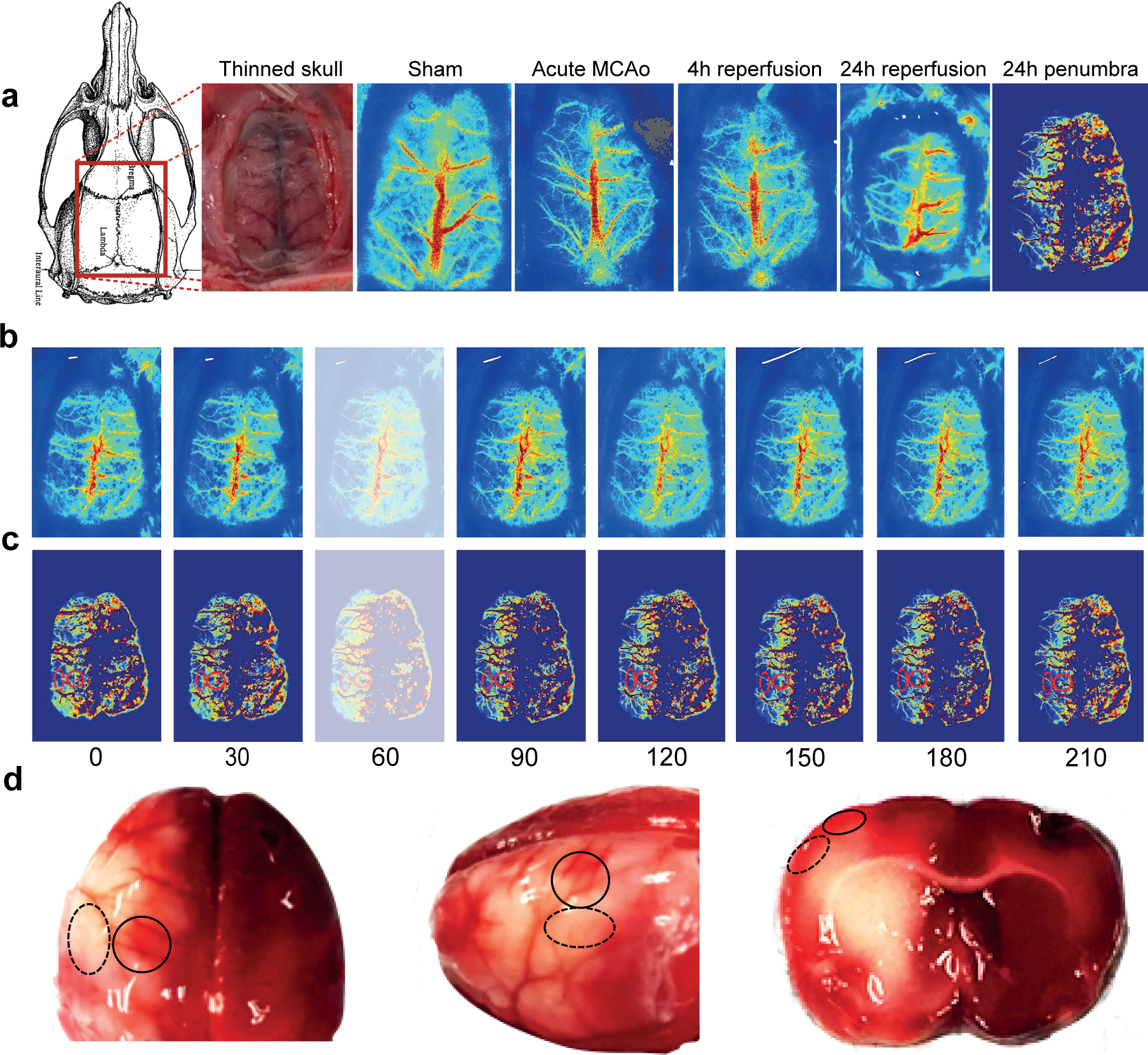
Laser speckle imaging reveals the fate of the penumbral tissue during the 4h experimental period. Following filament occlusion of the middle cerebral artery (MCAo) in Sprague-Dawley rats, a thin scull was made over the cortical area, where CBF was measured by the Laser Speckle Contrast Imaging (LSCI) (Fig.1a). Fig a. shows the LS signal in a sham animal, during acute MCAo, and 4 and 24h following reperfusion. We used the extent of LSCI signal reduction to subdivide hypoperfused cortex into penumbral calculated penumbral tissue^21-22,26-28^ after 4h of MACo and subsequent 24h of reperfusion. Row b shows the raw LS signal in a representative MACo animal during MCA occlusion at the tome points from 0-210 min at 30 min interval as where the 2PM measurements where performed. Row c shows the corresponding calculated penumbral tissue as it is affected by time after occlusion start. The fate of penumbral and core tissue was verified by TTC staining (row d). Thirty minutes after occlusion, an unexpected increase in blood flow in both ischemic and control animals was observed, resulting in elevated MTT and CTH values in both animal groups (Fig. 4a,b, Supplementary material Fig. 1b,c). These data are shown with transparent images, as it occurred in both MCAo animal and controls and seems to be a feature of our animal preparation, rather than of the stroke model.

**Supplementary figure 2.**
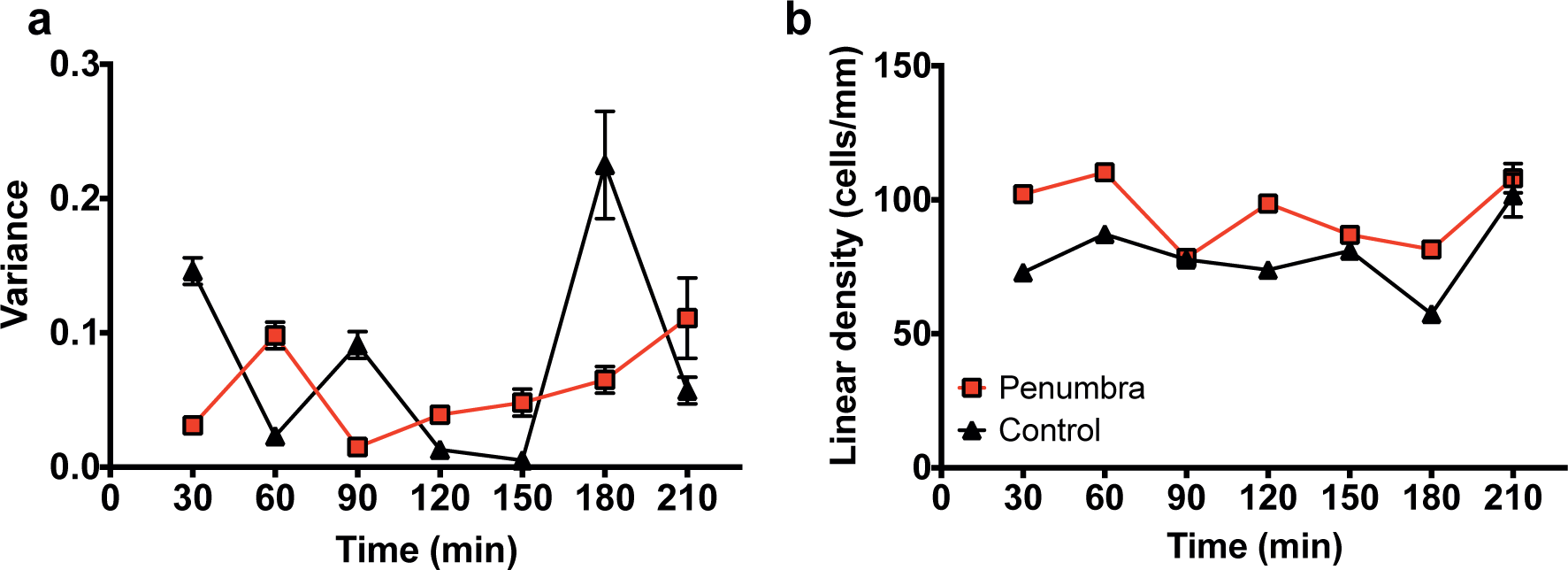
Penumbral RBC variance and linear density in MACo and control animals. TPM measurements of a) the variance of RBC velocities in the penumbral area, and b) linear density. Both measured parameters were not affected of MCAo or the fate of the penumbral tissue over time.

## Notes

### Competing Interest Statement

The authors have declared no competing interest.

